# Green Synthesis of Silver and Gold Nanoparticles by Aqueous Artemisia Pallens Extract

**DOI:** 10.1101/2024.04.17.589972

**Authors:** Ashutosh Kumar Verma

## Abstract

We successfully harnessed the potential of Artemisia pallens extracts for the eco-friendly biosynthesis of silver, gold, and silver-gold bimetallic nanoparticles, employing aqueous silver nitrate and chloroauric acid solutions. This innovative approach departs from traditional methods, often involving toxic chemical agents like hydrazine hydrate and sodium borohydride. In the quest for greener protocols, the biological route emerges as a non-toxic, straight-forward, and environmentally sound alternative, opening new avenues for translational research. This article discusses the production of silver, gold, and silver-gold nanoparticles using different species of Artemisia plants. Nanoparticle characterization was carried out using UV-visible spectrophotometry, TEM, XRD, and FTIR techniques. Microwave-assisted synthesis resulted in well-dispersed nanoparticles. In the case of silver nanoparticles, a spherical shape with a size of 6 nm was achieved using the microwave radiation-assisted method, while a size of 20 nm was obtained with UV-assisted synthesis. Gold nanoparticles exhibited diverse shapes, including spherical, triangular, prisms, trapezoids, and hexagonal, with a predominant size of 10 nm. The size range for gold nanoparticles varied from 10 nm to 400 nm.

## Introduction

In recent years, there has been a growing interest in the design and synthesis of noble metal nanoparticles with unique size and shape-dependent properties.^1^ Among these, silver nanoparticles (AgNPs), gold nanoparticles (AuNPs), and bimetallic nanoparticles (BMNPs) have emerged as promising candidates for a wide range of applications, including electronic and biomedical devices, environmental technologies, and energy technologies. ^2^ Biogenic synthesis methods,^3^ utilizing plant extracts as reducing agents,^4^ have gained attention as a green and sustainable approach for nanoparticle synthesis. The biogenic synthesis of Ag-NPs,^5^ AuNPs,^6^ and Ag-Au BMNPs^7^ has been explored using various plant extracts, such as Solidago canadensis, Stigmaphyllon ovatum, Gloriosa superba, pomegranate seeds, Arabic gum, and Moringa oleifera.^8–13^ Verma and Kumar provided a comprehensive review of the biosynthesis and applications of Ag and Au nanoparticles derived from Biological Systems.^14^ These biogenic synthesis methods offer control over the nanoparticles’ size, shape, and properties, opening up new possibilities for tailored applications.

The synthesis of AgNPs using biological methods is an easier and more benign alternative to chemical methods.^15^ AgNPs have unique properties, such as a large surface area, which make them suitable for various therapeutic applications. ^16,17^ Different parts of plant extracts have been utilized to synthesize AgNPs with silver ions as substrates. For example, extracts from Coffea arabica, olive leaf, Ocimum sanctum, and Combretum erythrophyllum have been used to produce spherical AgNPs with inhibitory effects against bacteria like E. coli and S. aureus.^18^ Additionally, the Mentha aquatica leaf extract and the ethyl acetate fraction of pomegranate leaves have been used for AgNP synthesis, resulting in smaller particles with enhanced antibacterial properties.^19^

Khali et al. synthesized spherical AgNPs using the aqueous extract of olive leaves, observing smaller particles at an alkaline pH of 8 compared to an acidic pH of 3.^20^ Dhand et al. demonstrated the synthesis of stable spherical AgNPs (20-30 nm) using the hydroalcoholic extract of Coffea arabica exposed to a silver nitrate solution.^21^ Smaller particle sizes were achieved with higher concentrations of silver nitrate (0.1 M) within 2 hours at room temperature, showing inhibitory effects against E. coli and S. aureus. Jain and Mehta employed the aqueous extract of Ocimum sanctum and its bioactive compound, quercetin, to synthesize spherical AgNPs.^19^ Quercetin yielded smaller particles (11.35 nm) with a narrow plasmon peak compared to the whole leaf extract of Ocimum sanctum (particle size: 14.6 nm), highlighting its significance. Padalia et al. utilized Tagetes erecta flowers’ aqueous extract to synthesize spherical and hexagonal AgNPs. ^22^ These AgNPs, when coupled with commercial antibiotics, showed enhanced antibacterial effects. Nouri et al. employed the aqueous extract of Mentha aquatica leaves to synthesize spherical AgNPs, with ultrasound enhancing the antibacterial activity and reducing the minimum inhibitory concentration compared to hydrothermal synthesis.^23^ Shaik et al. demonstrated that varying the volume of Origanum vulgare extract led to different-sized AgNPs, with increased extract volume resulting in smaller nanoparticles exhibiting bactericidal effects on both Gram-positive and Gram-negative bacteria. ^24^ Leela et al. investigated various plant leaf extracts, discovering that Helianthus annuus showed the most potent reduction of silver ions among the tested plants.^18^ Table 1 shows an analysis of the discussed plant-synthesized AgNPs.

**Table 1:**
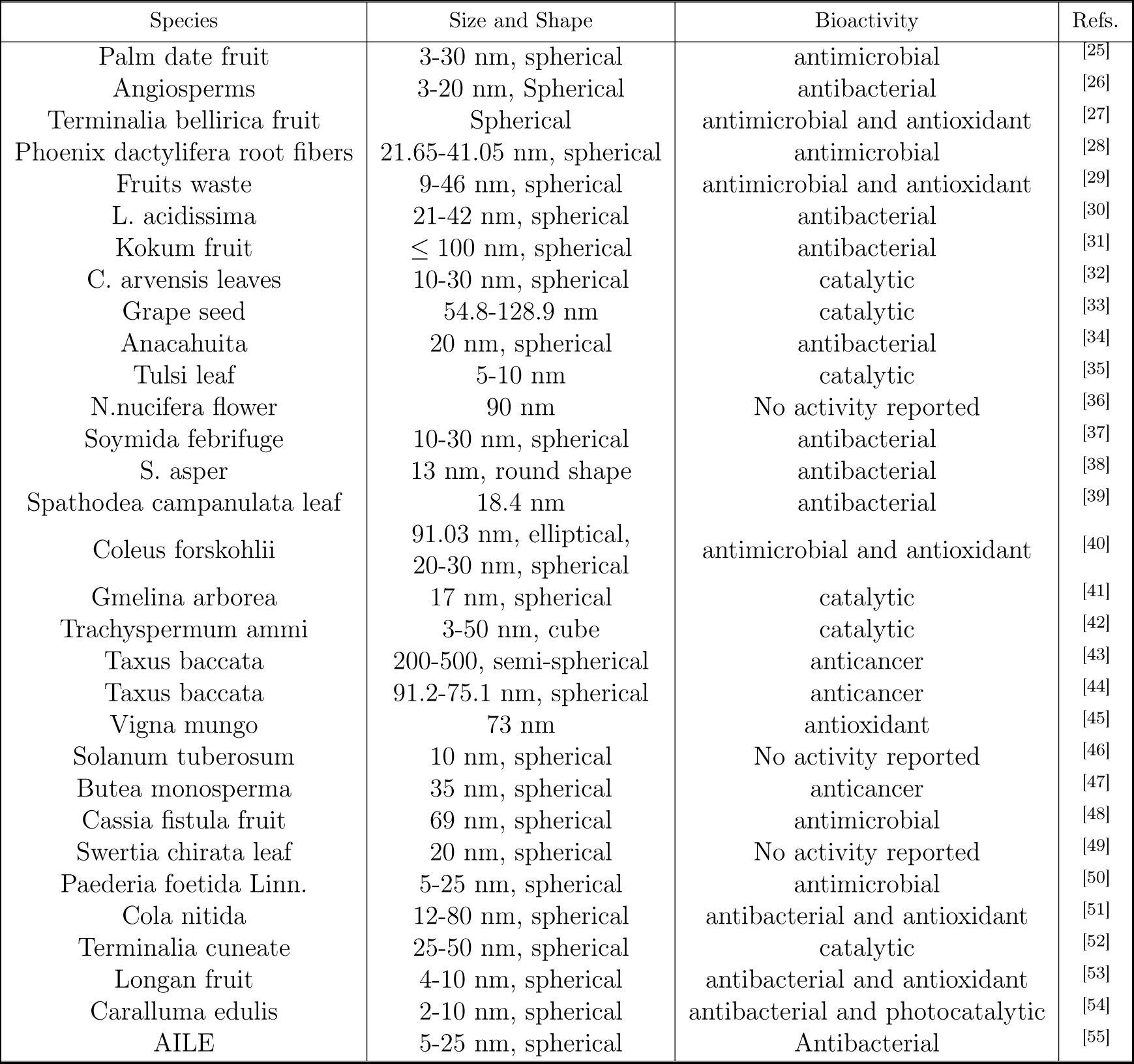
AgNPs from plants, size, shape, and biological activity.

AuNPs have diverse applications in fields such as gene therapy, catalysis, nanoelectronics, and disease diagnosis. ^2^ Green synthesis methods for AuNPs have gained attention due to concerns about the toxicity of chemical synthesis routes. Various plant extracts have been used for the biosynthesis of AuNPs. For example, Carica papaya and Catharanthus roseus leaf extracts were combined to produce predominantly spherical AuNPs with enhanced antibacterial effects.^56^ Synergizing AuNPs with antibiotics like rifampicin and kanamycin also resulted in spherical AuNPs with strong antibacterial properties.^57^ Other plant parts, including stems of Cannabis sativa, fruits of Amomum villosum and Pistacia atlantica, and aqueous extract of thyme, have been used for AuNP synthesis. ^35,58,59,59,60^ Mixtures of Olea europaea fruit extract and Acacia nilotica husk extract have shown significant antibacterial effects against specific bacteria. ^61^ Additionally, Croton caudatus Geisel leaf extract, Beta vulgaris pulp extract, Garcinia kola pulp, and Cryptolepis buchanani Roem aqueous extract have been utilized for the synthesis of stable and spherical AuNPs.^62–66^ The stability of the synthesized AuNPs has been confirmed through analysis of zeta potential. Green synthesis methods offer a safer and more sustainable approach to producing AuNPs with desirable properties. Table 2 shows an analysis of the discussed plant-synthesized AuNPs.

**Table 2:**
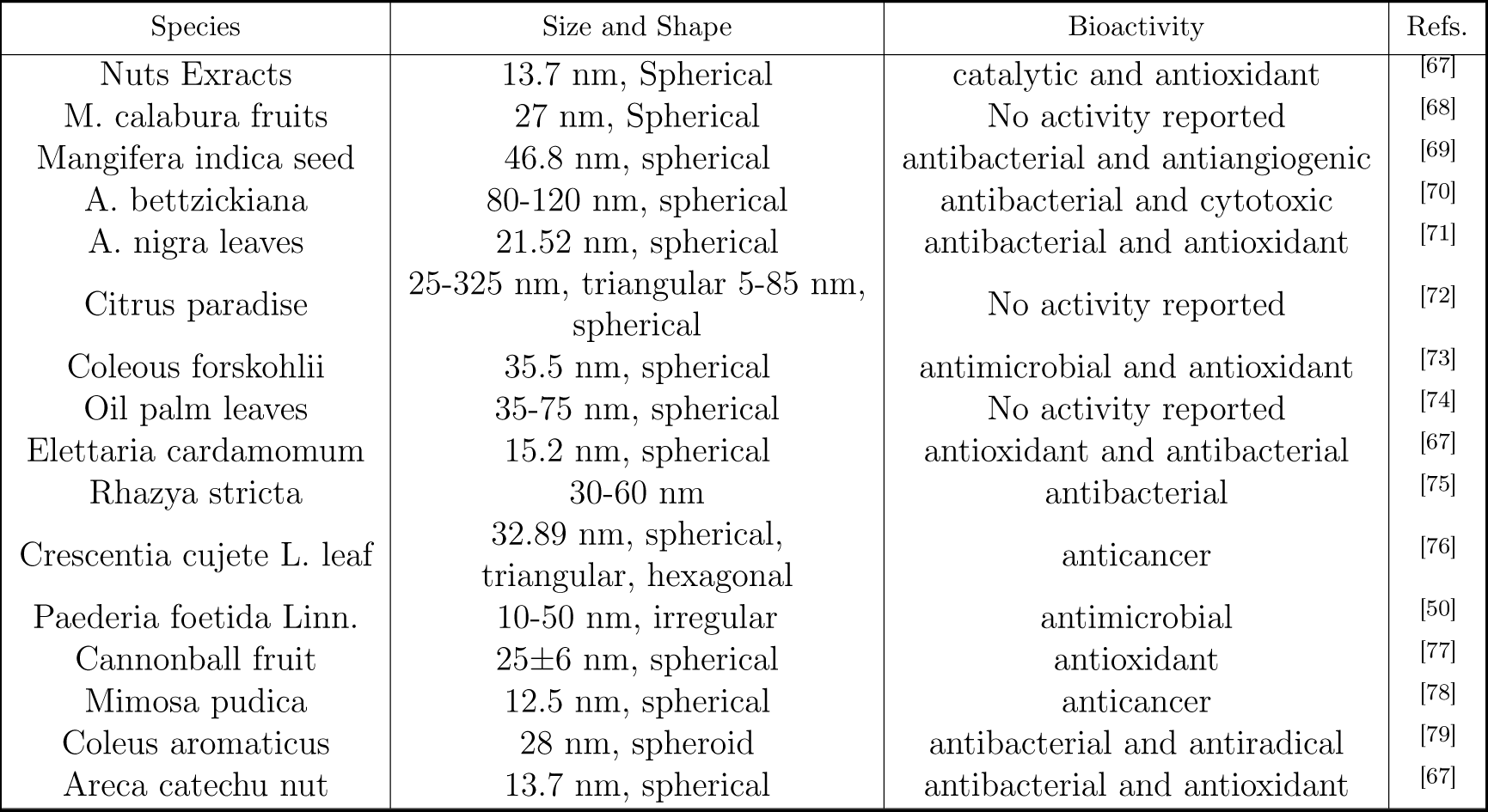
AuNPs from plants, size, shape, and biological activity.

| Species | Size and Shape | Bioactivity | Refs. |
| --- | --- | --- | --- |
| Nuts Extracts | 13.7 nm, Spherical | catalytic and antioxidant | [67] |
| *M. calabura* fruits | 27 nm, Spherical | No activity reported | [68] |
| *Mangifera indica* seed | 46.8 nm, spherical | antibacterial and antiangiogenic | [69] |
| *A. bettzickiana* | 80-120 nm, spherical | antibacterial and cytotoxic | [70] |
| *A. nigra* leaves | 21.52 nm, spherical | antibacterial and antioxidant | [71] |
| Citrus paradise | 25-325 nm, triangular 5-85 nm, spherical | No activity reported | [72] |
| *Coleous forskohlii* | 35.5 nm, spherical | antimicrobial and antioxidant | [73] |
| Oil palm leaves | 35-75 nm, spherical | No activity reported | [74] |
| *Elettaria cardamomum* | 15.2 nm, spherical | antioxidant and antibacterial | [67] |
| *Rhazya stricta* | 30-60 nm | antibacterial | [75] |
| *Crescentia cujete* L. leaf | 32.89 nm, spherical, triangular, hexagonal | anticancer | [76] |
| *Paederia foetida* Linn. | 10-50 nm, irregular | antimicrobial | [50] |
| Cannonball fruit | 25±6 nm, spherical | antioxidant | [77] |
| *Mimosa pudica* | 12.5 nm, spherical | anticancer | [78] |
| *Coleus aromaticus* | 28 nm, spheroid | antibacterial and antiradical | [79] |
| *Areca catechu* nut | 13.7 nm, spherical | antibacterial and antioxidant | [67] |

BMNPs offer improved electronic, optical, and catalytic properties compared to monometallic nanoparticles.^80^ They can exist in different forms, such as alloys, core-shell structures, and contact aggregates (see Table 3). Ag-Au BMNPs have shown advancements in drug delivery and nanomedicine. ^81^ Various plant extracts have been used for the synthesis of Ag-Au BMNPs, resulting in different shapes and properties. For example, the aqueous extract of Solidago canadensis and the leaf extract of Stigmaphyllon ovatum has been used to synthesize Ag-Au alloys with similar shapes to gold nanoparticles. ^9^ Gloriosa superba extract has been employed for the synthesis of Ag-Au nanoalloys with enhanced antibacterial activity.^10^ Pomegranate seed juice and Arabic gum have also been used for the green synthesis of Ag-Au BMNPs, with reaction conditions influencing the size and dispersion of the nanoparticles.^11^ Additionally, Moringa oleifera leaf extract has been utilized for the bio-fabrication of stable Ag-Au nanoalloys.^13^ The properties and applications of BMNPs depend on their size, shape, and metal composition. Synthetic strategies for Ag-Au BMNPs include continuous growth, seed-mediated growth, and galvanic replacement reactions. These strategies allow for control over the growth and formation of alloyed, core-shell, or hetero-structured BMNPs. Ligands and reaction parameters play important roles in shaping the morphology of the BMNPs during synthesis. The design and synthesis of noble metal nanostructures, including Ag-Au BMNPs, have gained significant interest due to their unique properties and potential applications in various fields.

**Table 3:**
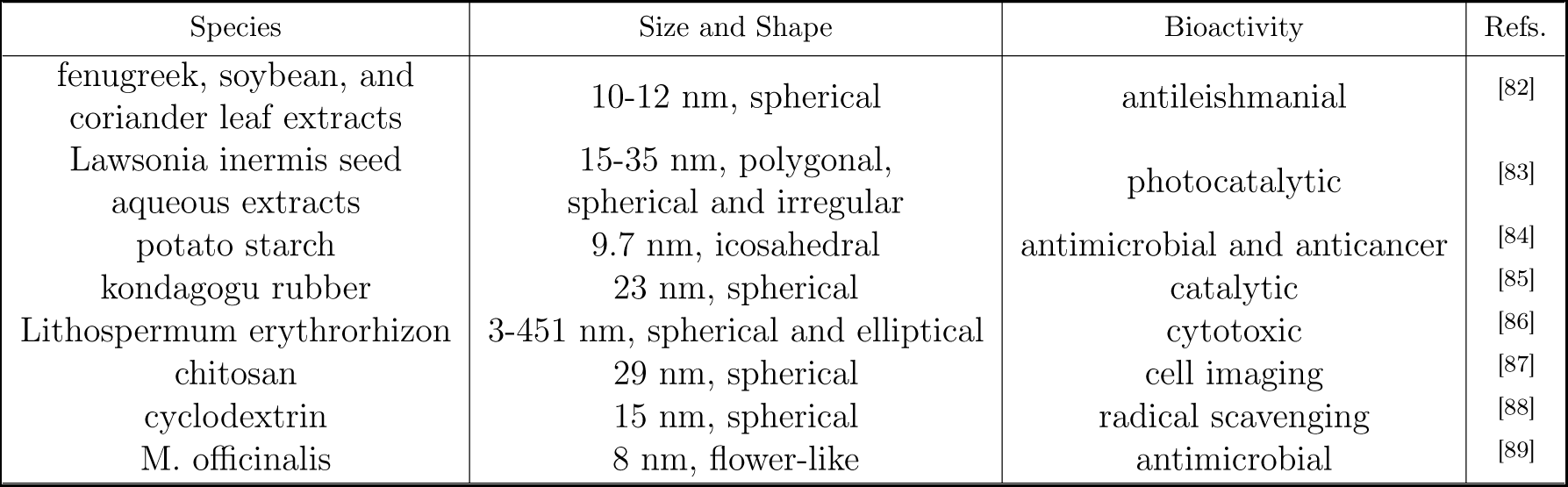
Ag-Au alloy and core-shell nanoparticles from plants, size, shape, and biological activity.

The use of Artemisia pallens for AgNPs was first reported by Arde et al.^90^ In this work, we synthesized AgNPs, AuNPs, and Ag-Au bimetallic NPs using Artemisia pallens, although other Artemisia species have been utilized in the past. Developing environmentally friendly processes for synthesizing nanoscale materials is crucial, and Ag-Au BMNPs have potential applications in various fields due to their unique properties. By delving deeper into their synthesis, characterization, and functional properties, this research paper contributes to the growing body of knowledge on biogenic synthesis approaches and provides valuable insights for future advancements in the field of nanotechnology.

## Materials and Methods

### Chemical and reagents

The Artemisia plant was obtained from the local market in Chennai, India. All reagents used in the experiment were of analytical grade, including Silver Nitrate (AgNO_3_) and Gold Chloride (HAuCl_4_.3H_2_O) purchased from Sigma-Aldrich, India. Distilled water was utilized throughout the experiment.

### Preparation of plant extract

The Artemisia plant underwent multiple rinses with distilled water before being finely ground with a mortar and pestle. Subsequently, 40 g of the ground material was combined with 100 mL of deionized water, gently stirred at 30 °C for 1 minute, and filtered to obtain an aqueous extract. Separate extracts were prepared from various components of the Artemisia plant, including the stem, leaves, and entire plant.

### Extraction of nanoparticles from the plant material and their characterization

Nanoparticle extraction typically involves room-temperature material crushing, followed by water extraction. To validate nanoparticle synthesis, standard characterization methods such as UV-VIS, DSL, ZP, FTIR, XRD, EDS, AES, SPM, XPS, STM, AFM, TEM, and SEM are employed. Initially, visual observations are used to study nanoparticle formation, followed by UV-visible spectrum analysis. High-resolution transmission electron microscopy (HR-TEM) is employed to determine morphology (size and shape). The element content of the nanoparticle preparation is obtained through X-ray studies.

### AgNPs synthesis using Artemisia Plant

AgNPs possess conductive, optical, ^25^ and diagnostic properties, making them highly versatile. Their bio-production has garnered significant attention due to their remarkable antibacterial^67,91^ capabilities, particularly in controlling multi-drug resistant microorganisms. Additionally, these nanoparticles exhibit noteworthy anti-cancer^43,44,47,67,92^ effects. In this study, we primarily characterized the formation of nanoparticles by analyzing their UV-visible absorption spectrum. When subjected to electromagnetic radiation of a specific wavelength, the molecules undergo electronic transitions. Our UV-visible spectrophotometer measured the surface plasma resonance of AgNPs at a wavelength near 450 nm (Fig. 1, 3, 5). The size of the synthesized AgNPs (see Table 4) was estimated from TEM images (Fig. 7).

**Table 4:**
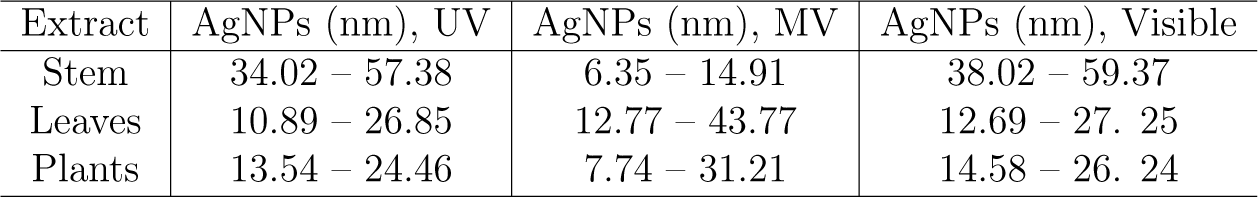
Size of the AgNPs under visible, UV, and MV-assisted synthesis.

### AuNPs synthesis using Artemisia Plant

AuNPs possess catalytic, antibacterial, antioxidant, and anticancer properties. ^67^ The bio-production of these nanoparticles has garnered significant attention due to their antibacterial potential,^69,70,91^ which may aid in combating multi-drug resistant microorganisms. Additionally, they have demonstrated anti-cancer effects. ^43,44,47^ In this study, nanoparticle formation was primarily characterized using UV-visible absorption spectroscopy, which measures the electronic transitions of molecules upon exposure to electromagnetic radiation of specific wavelengths. The AuNPs exhibited surface plasma resonance at approximately 550 nm, as determined by a UV-visible spectrophotometer (Fig. 2, 4, 6). The size of the synthesized AuNPs(see Tables 5-7) was estimated from TEM images(Fig. 8).

**Table 5:** Size of the AuNPs using leaves extract (visible, UV, and MV-assisted synthesis).

| Shape | AuNPs (nm), UV | AuNPs (nm), MV | AuNPs (nm), Visible |
| --- | --- | --- | --- |
| Triangular | 51.39 – 467.16 | 9.49 – 51.33 | 44.87 – 643.28 |
| Hexagonal | 17.99 – 53.28 | 12.42 – 47.50 | 74.70 – 507.46 |
| Pentagonal |  | 27.64 – 38.04 | 72.92 – 73.51 |
| Truncated Triangular | 51.39 – 467.16 | 29.73 – 108.24 | 422.15 – 643.20 |

**Table 6:** Size of the AuNPs using stem extract (visible, UV, and MV-assisted synthesis).

| Shape | AuNPs (nm), UV | AuNPs (nm), MV | AuNPs (nm), Visible |
| --- | --- | --- | --- |
| Triangular | 30.98 -135.84 | 53.87 – 203.27 | 224.34 – 327.5 |
| Hexagonal | 19.33 – 74.80 | 34.47 – 156.74 | 212.95 – 437.92 |
| Truncated Triangular | 67.48 – 214.45 | 109.32 – 172.35 |  |

**Table 7:**
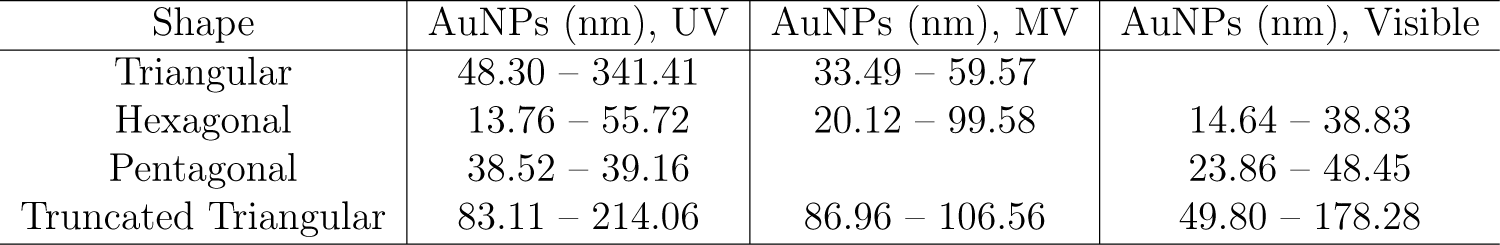
Size of the AuNPs using plant extract (visible, UV, and MV-assisted synthesis).

## Results and Discussion

### Using plant extract

The formation of Ag and Au NPs was studied using an extract from the Artemisia plant in a novel medium. Different concentrations of AgNO_3_ solution (ranging from 1 mM to 10 mM) were mixed with the plant extract in a ratio of 9:1. The absorption spectrum for Ag ion concentrations exhibited peak profiles at approximately 450 nm (Fig. 1). Similarly, HAuCl_4_.3H_2_O solution (ranging from 0.1 mM to 1 mM) was mixed with the plant extract in the same ratio. The absorption spectrum for Au ion concentrations displayed peak profiles around 550 nm (Fig. 2).

**Figure 1:**
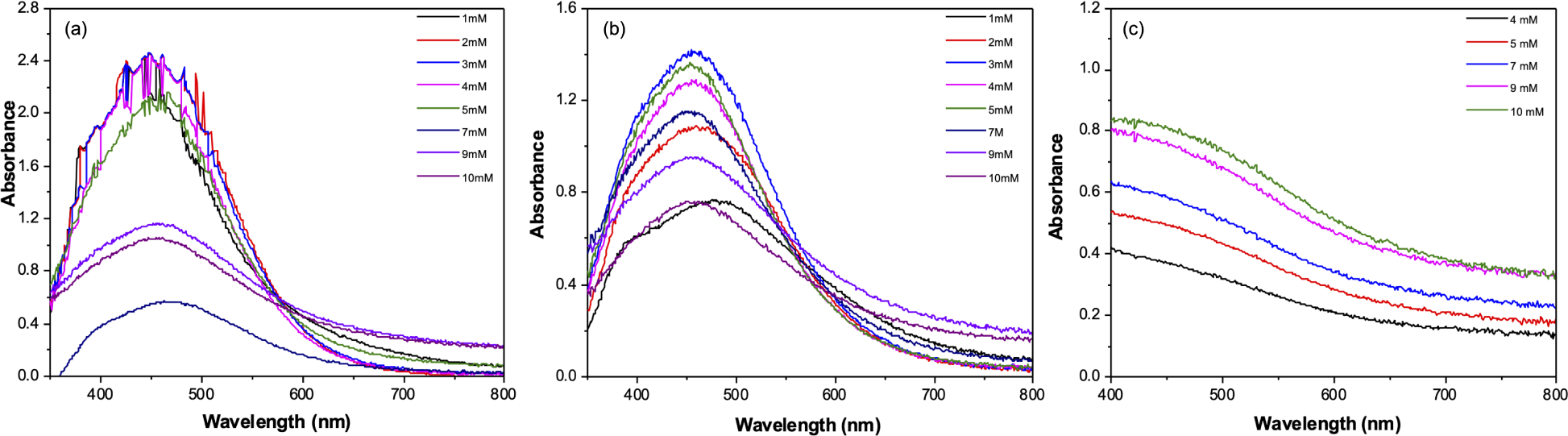
Absorption spectrum of formation of AgNPs in the presence of 1, 2, 3, 4, 5, 7, 9, 10 mM AgNO_3_ at 24h reaction time under (a) visible radiation, (b) UV radiation and (c) less than 1 minutes reaction time under microwave radiation.

**Figure 2:**
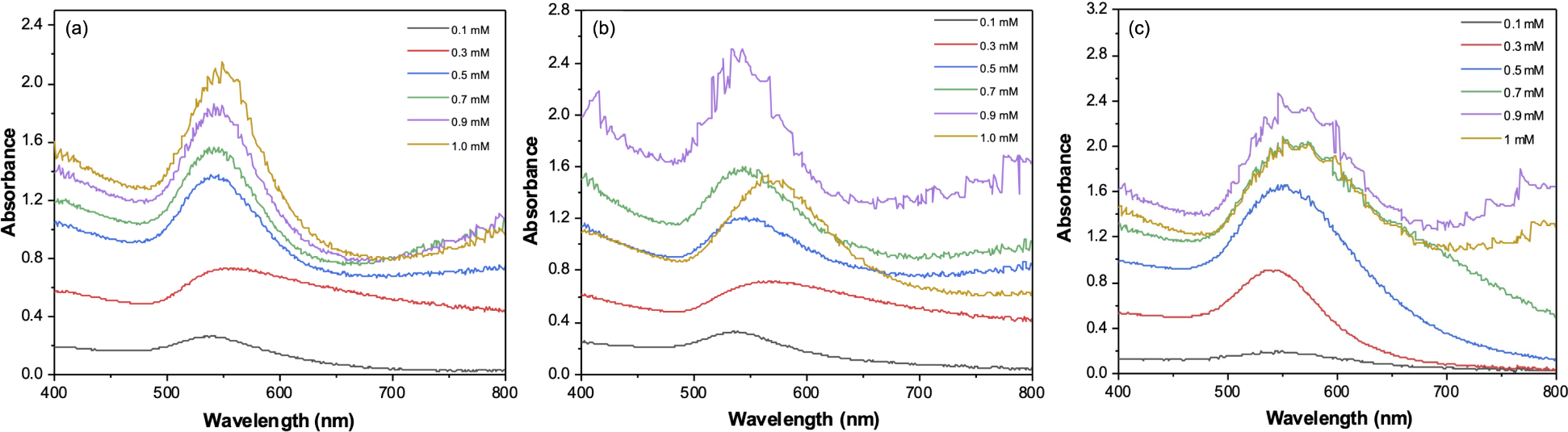
Absorption spectrum of formation of AuNPs in the presence of 0.1, 0.3, 0.5, 0.7, 0.9, 1 mM HAuCl_4_.3H_2_O at 6h reaction time under (a) visible radiation, (b) UV radiation and (c) less than 1 minutes reaction time under microwave radiation.

### Using stem extract

The formation of Ag and Au NPs using Artemisia stem extract was studied using different media, including microwave, UV, and visible radiation. AgNO_3_ solutions ranging from 1 mM to 10 mM were mixed with stem extract in a ratio of 9:1. The absorption spectrum of the Ag ion concentrations exhibited peaks at approximately 450 nm (Fig. 3). Similarly, HAuCl_4_.3H_2_O solutions ranging from 0.1 mM to 1 mM were mixed with stem extract in the same ratio. The absorption spectrum of the Au ion concentrations displayed peaks around 550 nm (Fig. 4).

**Figure 3:**
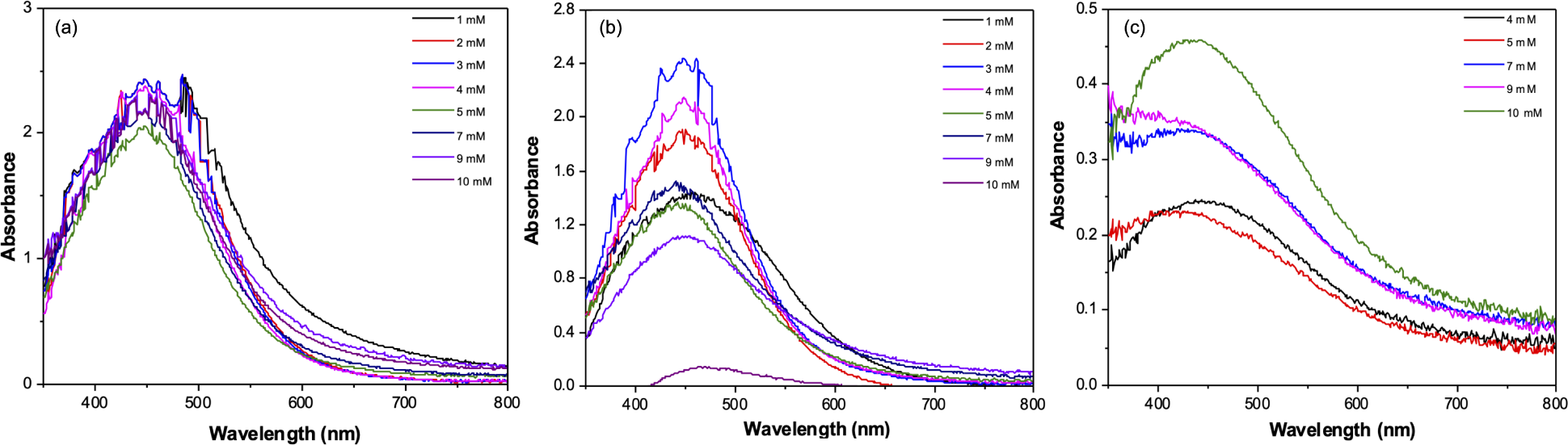
Absorption spectrum of formation of AgNPs in the presence of 1, 2, 3, 4, 5, 7, 9, 10 mM AgNO_3_ at 24h reaction time under (a) visible radiation, (b) UV radiation and (c) less than 1 minutes reaction time under microwave radiation.

**Figure 4:**
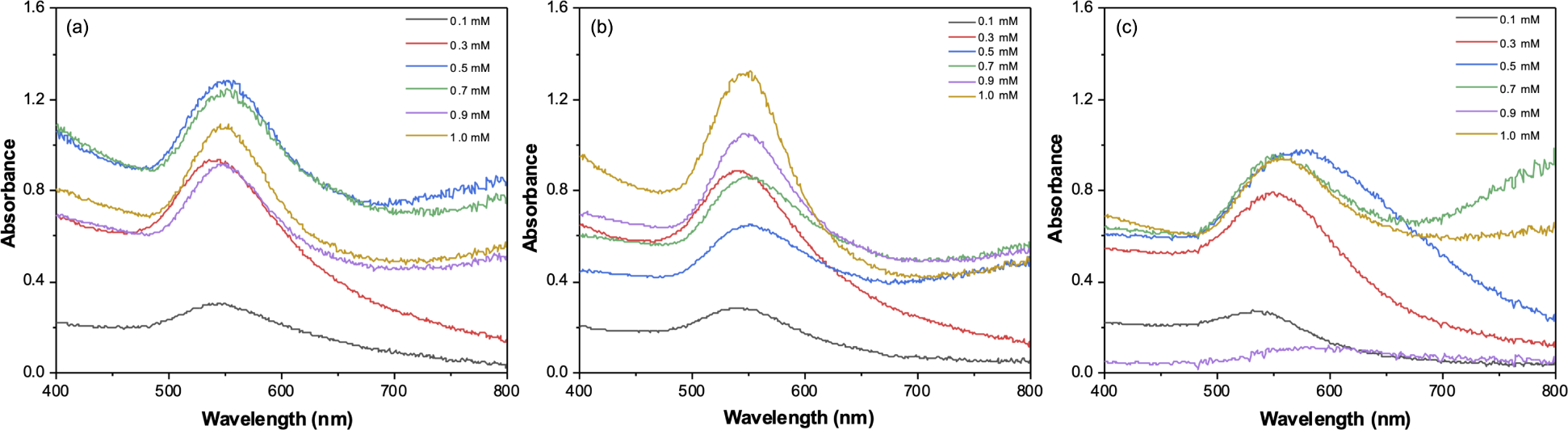
Absorption spectrum of formation of AuNPs in the presence of 0.1, 0.3, 0.5, 0.7, 0.9, 1 mM HAuCl_4_.3H_2_O at 6h reaction time under (a) visible radiation, (b) UV radiation and (c) less than 1 minutes reaction time under microwave radiation.

### Using leaves extract

The formation of Ag and Au NPs using Artemisia pallens was studied using three different media. AgNO_3_ solutions ranging from 1 mM to 10 mM were mixed with leaf extract in a 9:1 ratio. The absorption spectrum of Ag ion concentrations exhibited profiles with a peak at approximately 450 nm (Fig. 5). Similarly, HAuCl_4_.3H_2_O solutions ranging from 0.1 mM to 1 mM were mixed with leaves extract in a 9:1 ratio. The absorption spectrum of Au ion concentrations displayed profiles with a peak at around 550 nm (Fig. 6).

**Figure 5:**
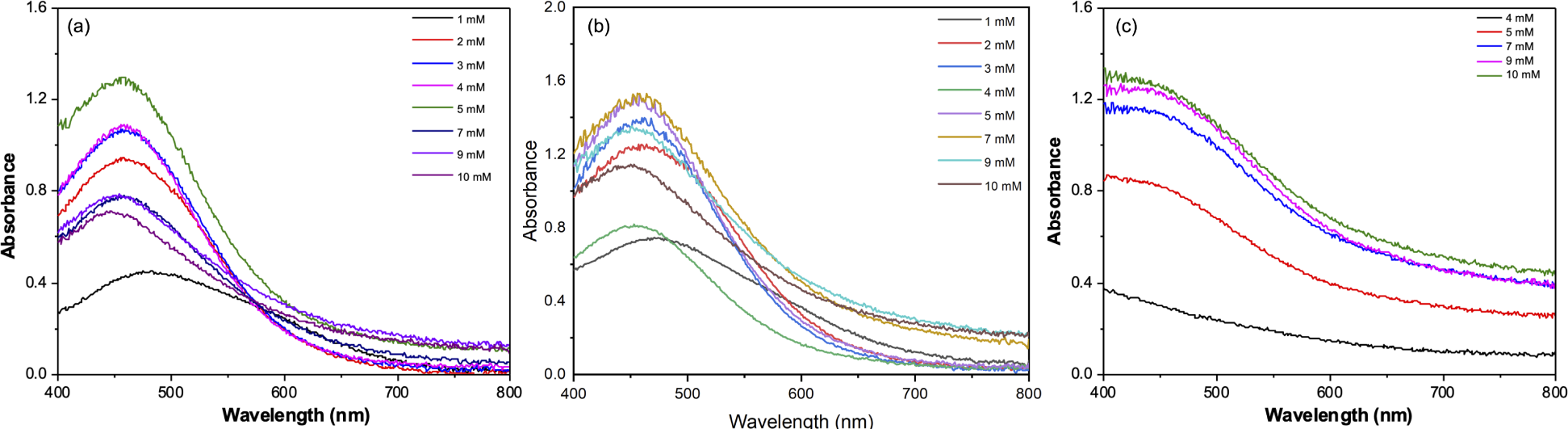
Absorption spectrum of formation of AgNPs in the presence of 1, 2, 3, 4, 5, 7, 9, 10 mM AgNO_3_ at 24h reaction time under (a) visible radiation, (b) UV radiation and (c) less than 1 minutes reaction time under microwave radiation.

**Figure 6:**
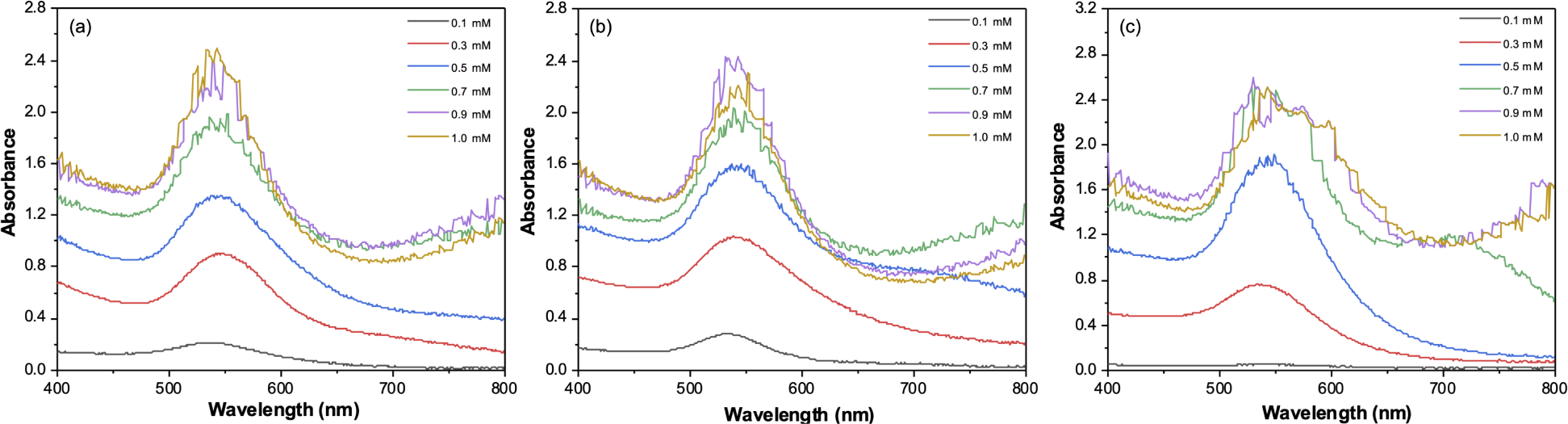
Absorption spectrum of formation of AuNPs in the presence of 0.1, 0.3, 0.5, 0.7, 0.9, 1 mM HAuCl_4_.3H_2_O at 6h reaction time under (a) visible radiation, (b) UV radiation and (c) less than 1 minutes reaction time under microwave radiation.

### TEM analysis of AgNPs, AuNPs, and Ag-Au BMNPs

TEM analysis is a powerful technique for characterizing the size, shape, and morphology of nanoparticles, particularly AgNPs and AuNPs synthesized from Artemisia pallens. It provides valuable insights into the structural properties of these nanoparticles. By analyzing the AgNPs and AuNPs synthesized from Artemisia pallens, TEM analysis helps understand the synthesis method’s effectiveness and facilitates further optimization if necessary. The analysis revealed a high density of mostly spherical AgNPs (see Fig. 7(a)), with many measuring approximately 6.35 nm or larger (see Table 4). Additionally, some nano-aggregates were observed (see Fig. 7(b)). These findings align with previous studies using extracts from field-grown parts of various species (see Table 1, 2, and 3). Alloy and core-shell type BMNPs were also synthesized with varying sizes of between 6.00 to 200.00 nm (Table 8).

**Figure 7:**
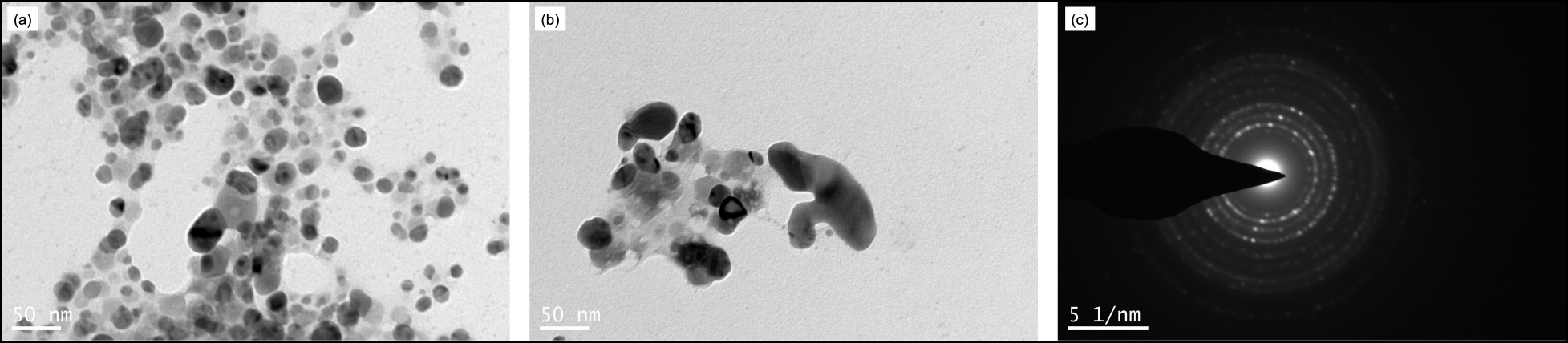
TEM images of Microwave-assisted AgNPs synthesized using Artemisia plant extract (a) a high density of spherical AgNPs, (b) nano-aggregates, and (c) SAED

**Table 8:** Size of the AuNPs using plant extract (visible, UV, and MV-assisted synthesis).

| BMNPs | Size (nm), UV | Size (nm), MV | Size (nm), Visible |
| --- | --- | --- | --- |
| Core-shell | 10.00-200.00 | 6.30-50.00 | 20.00-100.00 |
| Alloy |  | 6.00 – 50.00 |  |

The SAED patterns in Fig. 7(c) and Fig. 8(c) confirm the presence of pure silver and gold in the AgNPs and AuNPs, respectively. Additionally, EDAX studies indicated the presence of a characteristic absorption peak at 22 keV (see Fig. 9(a)) and this confirmed the presence of Ag atoms in the AgNPs. EDAX analysis also confirmed the occurrence of Au atoms in the AuNPs since a characteristic absorption peak was seen at 10 keV indicating the presence of pure Au with no other contaminants (see Fig. 9(b)). TEM images of the AuNPs formed by the Artemisia pallens also revealed that the AuNPs were well-dispersed. A higher density and an immense diversity was observed in the geometry and size of the AuNPs (see Fig. 8(a)) obtained with the Artemisia pallens unlike the case of microwave-assisted synthesis with which only particles of much smaller size range were obtained (see Table 5). A mixture of AuNPs of diverse shapes including spherical, triangular, truncated triangular, pentagonal, and hexagonal was obtained with a predominant size of 9.49 nm. The size of AuNPs was between 9.49 nm and 643.2 nm. Similar shapes of nanoparticles have been commonly obtained with extracts of field-grown parts of many species(see Table 5-7).

**Figure 8:**
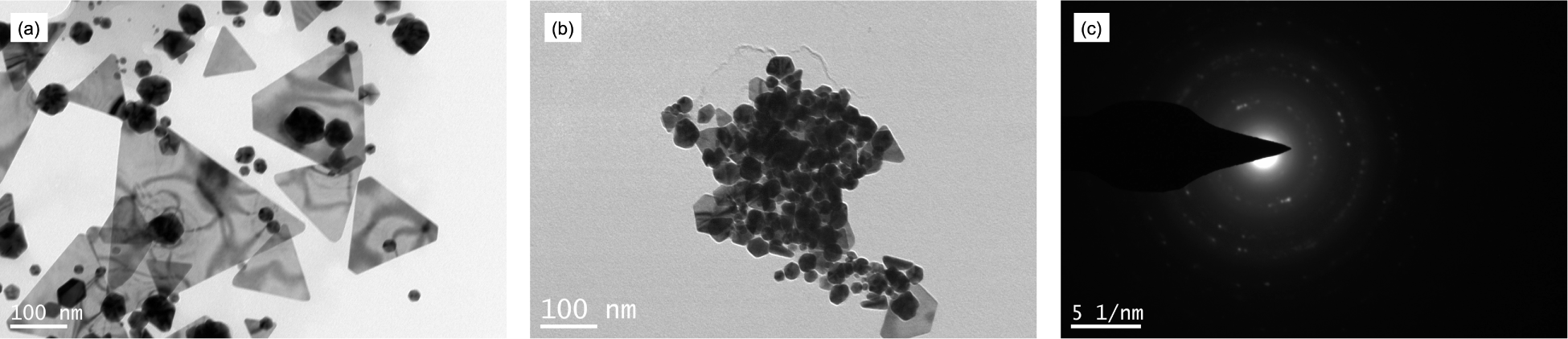
TEM images of Microwave-assisted AuNPs synthesized using Artemisia plant extract (a) a high density of spherical AgNPs, (b) nano-aggregates, and (c) SAED

**Figure 9:**
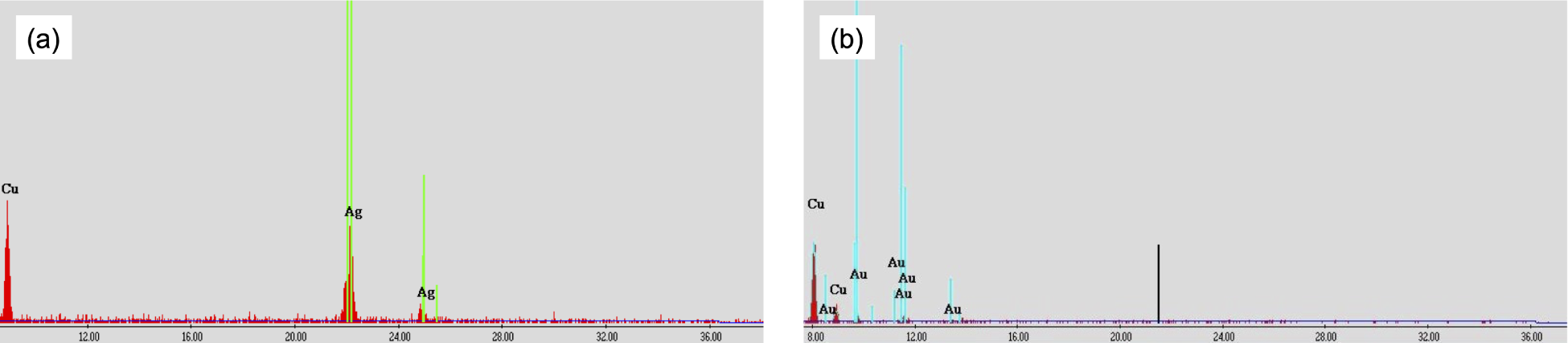
EDAX analysis was performed on well-dispersed nanoparticles (a) Ag nanoparticles, (b) Au nanoparticles.

Additionally, some nano-aggregates were observed (see Fig. 8(b)).

### XRD analysis of AgNPs and AuNPs

The distinct diffraction at 38.2°, 41.8°,46.2° 77.5° and 86° are indexed to the [111], [200], [220], [311], and [222] crystallographic planes of AgNPs, respectively which indicates crystalline nature of AgNPs (see Fig. 10(a)). The observed set of lattice planes was indexed based on the face-centered cubic (FCC) structure of AgNPs by comparing the XRD data with JCPD Card No. 89-3722.

**Figure 10:**
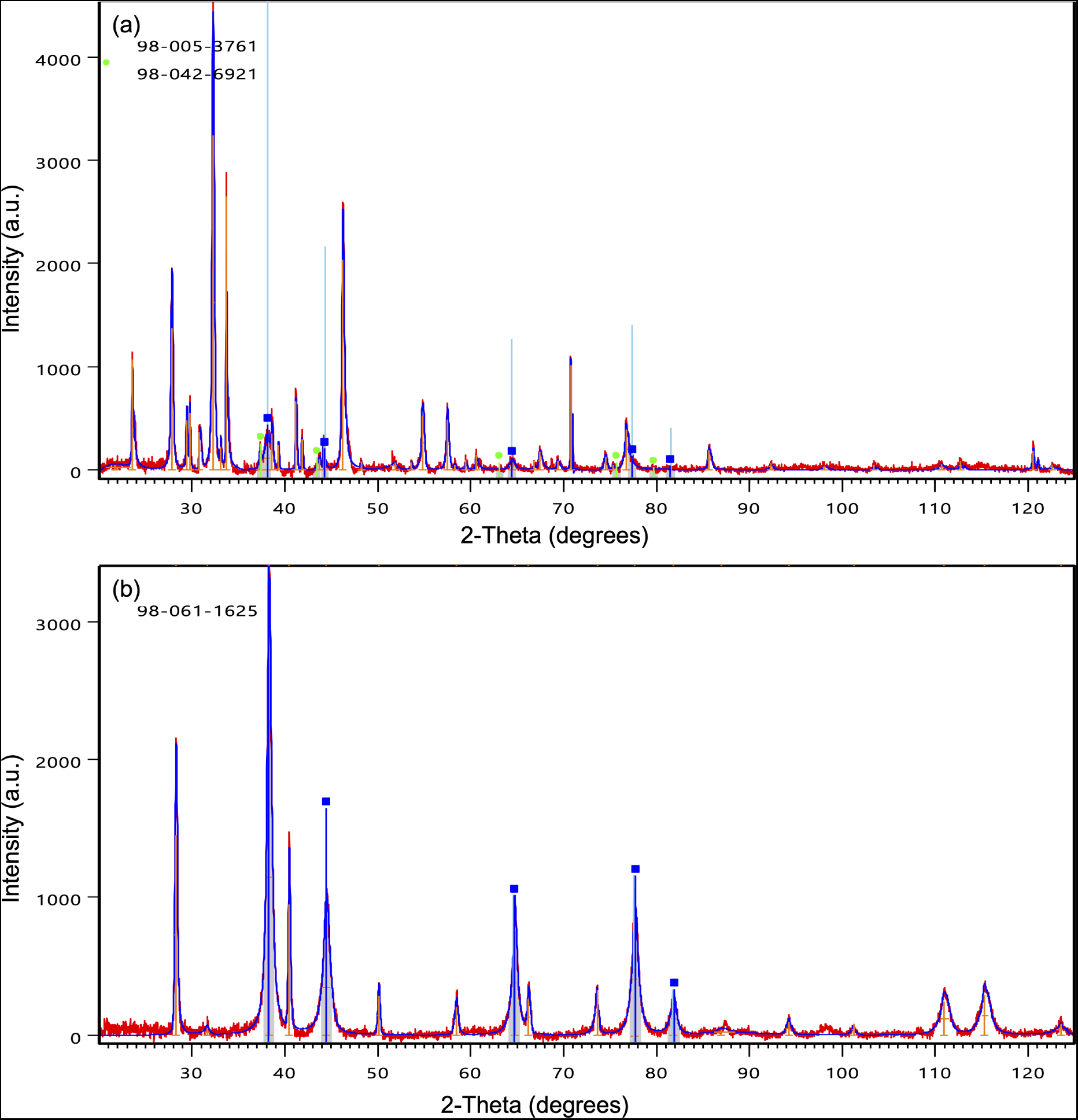
XRD analysis of nanoparticles synthesized at 30 *◦C* by Artemisia plant extract (a) Ag nanoparticles, (b) Au nanoparticles.

**Figure 11:**
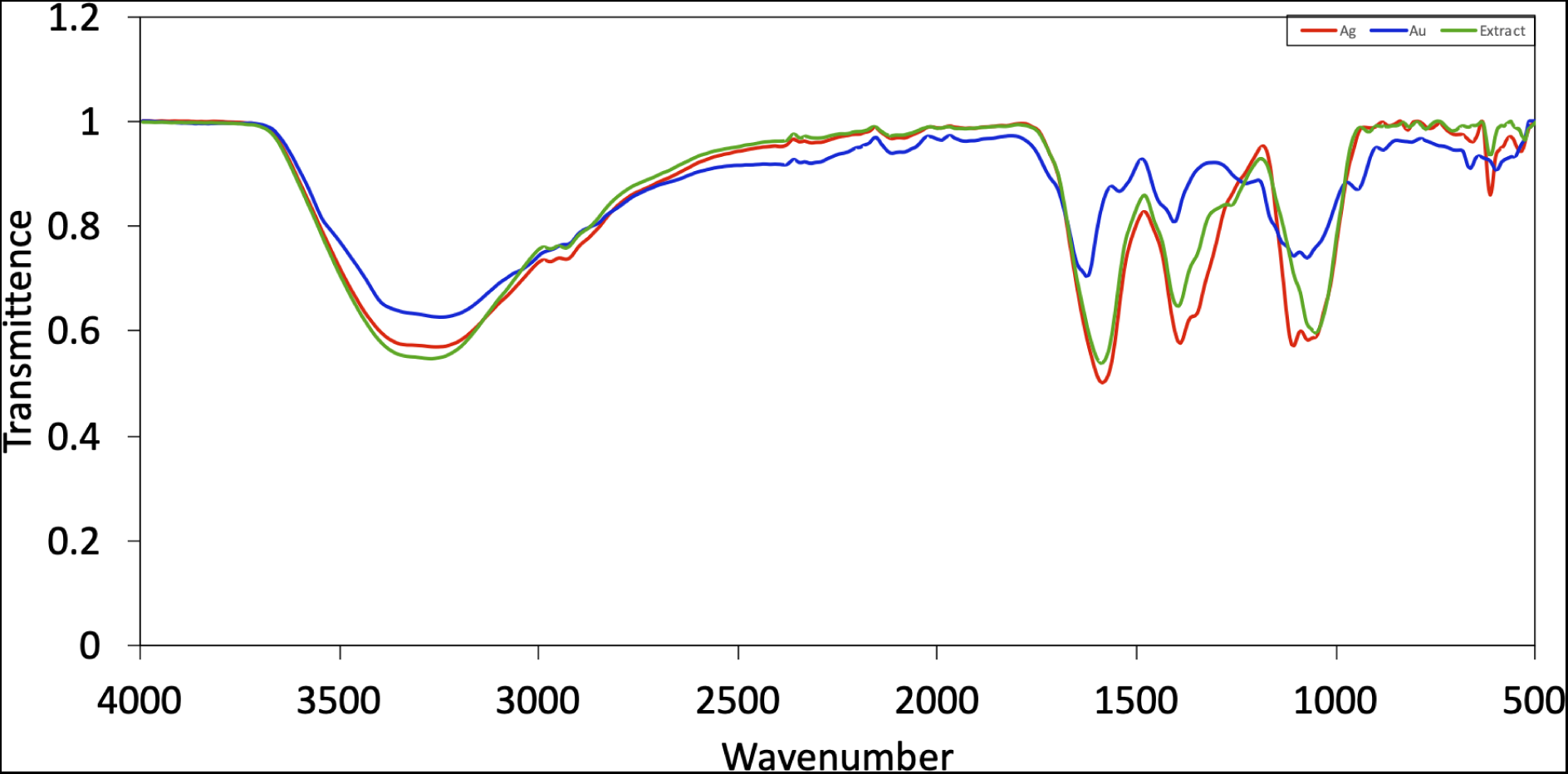
FTIR analysis of extract and the Ag and Au NPs synthesized by Artemisia plant extract

The unpredicted crystalline at other peaks is also present this could be due to the organic compound belonging to the leaf extract. The typical x-ray diffractograms of the Au NPs had 2*θ* values at 38.25, 46, 64.55, and 77.49 that can be indexed to (111, 200, 220) and (311) facets of faced centered cubic structure (see Fig. 10(b)). Other picks of 2*θ* values might have resulted from the crystallization of some bio-organic compound or phenolic compound of the extract on the surface of the Au NPs.

### FTIR analysis of AgNPs and AuNPs

Figure 11 displays the FTIR spectra of the Artemisia extract, AgNPs, and AuNPs. The FTIR spectrum exhibits absorbance peaks at wave numbers 3278, 1642, 1585, 1393, 1344, 1105, 1048, 1011, and 614. A common band at 1393.05 cm*^−^*^1^ is observed for the extract, AgNPs, and AuNPs, potentially corresponding to dimethyl or nitro compounds. The intense broad peak at 3278 cm-1 is typical of the hydroxyl functional group of phenols and alcohols. Additionally, the bands at 1442 and 1585 cm*^−^*^1^ may correspond to amides or primary amines, while the bands at 1344, 1393, 1105, 1048, and 1011 cm*^−^*^1^ indicate O-H bending and C-O stretching vibrations of phenols. Therefore, the FTIR studies suggest that bio-reduction and phytocapping of AgNPs and AuNPs by phytochemicals in the Artemisia extract likely played a significant role in their formation.

### Antimicrobial Activity

AgNPs are extensively researched for their antimicrobial properties, notably their potential antibacterial effects (see Table **??**). Arde et al. assessed the antimicrobial activity of AgNPs synthesized using Artemisia pallens plant extracts. They evaluated antimicrobial activity against pathogenic bacteria using a 5mm diameter paper disc method with 0.05 mg/mL samples. These AgNPs demonstrated superior antimicrobial effectiveness against Staphylococcus aureus, Bacillus cereus, and Escherichia coli compared to antibiotics. Chloramphenicol, at a concentration of 0.5 mg/mL, served as the control antimicrobial agent.^90^ Therefore, we did not investigate the antimicrobial activity of Ag and Au NPs in this study.

## Conclusion

In conclusion, we have demonstrated the use of a natural, renewable, non-toxic, and low-cost aqueous extract of Artemisia pallens as a bio-reducing agent for the effective synthesis of AgNPs, AuNPs, and Ag-Au BMNPs with radiation under visible, UV and microwave. Different parts of Artemisia, such as stem, leaves, and entire plant extract, were studied as reducing agents for the synthesis of NPs. The potential for NPs synthesis varies in different organs of the same species of plants. The plant enabled the synthesis of well-dispersed AgNPS, AuNPs, and Ag-Au BMNPs of different sizes and shapes. AgNPs of spherical shape with a size of 6 nm were found in the microwave radiation-assisted method, whereas, the size of the AgNPs was 20 nm in UV-assisted synthesis. AuNPs of diverse shapes, such as spherical, triangular, prisms, trapezoids, and hexagonal were obtained with the predominant size of 10 nm. The size of AuNPs ranged from 10 nm to 400 nm.

## Acknowledgments

We gratefully acknowledge the Sophisticated Analytical Instrument Facility at IIT Madras for FTIR analysis. Additionally, we extend our thanks to the Department of Metallurgical and Materials Engineering, IIT Madras, for providing access to the Transmission Microscopy Facility and Central X-ray diffraction facility, enabling TEM measurements and XRD analysis.

